# NeuronUnit: A package for data-driven validation of neuron models using SciUnit

**DOI:** 10.1101/665331

**Authors:** Richard C. Gerkin, Justas Birgiolas, Russell J. Jarvis, Cyrus Omar, Sharon M. Crook

## Abstract

Validating a quantitative scientific model requires comparing its predictions against many experimental observations, ideally from many labs, using transparent, robust, statistical comparisons. Unfortunately, in rapidly-growing fields like neuroscience, this is becoming increasingly untenable, even for the most conscientious scientists. Thus the merits and limitations of existing models, or whether a new model is an improvement on the state-of-the-art, is often unclear.

Software engineers seeking to verify, validate and contribute to a complex software project rely on suites of simple executable tests, called “unit tests”. Drawing inspiration from this practice, we previously developed *SciUnit*, an easy-to-use framework for developing data-driven “model validation tests” – executable functions, here written in Python. Each such test generates and statistically validates predictions from a model against one relevant feature of empirical data to produce a score indicating agreement between the model and the data. Suites of such validation tests can be used to clearly identify the merits and limitations of existing models and developmental progress on new models.

Here we describe *NeuronUnit*, a library that builds upon *SciUnit* and integrates with several existing neuroinformatics resources to support the validation of single-neuron models using data gathered by neurophysiologists and neuroanatomists. *NeuronUnit* integrates with existing technologies like Jupyter, Pandas, NeuroML and resources such as NeuroElectro, The Allen Institute, and The Human Brain Project in order to make neuron model validation as easy as possible for computational neuroscientists.

## 1 INTRODUCTION

### 1.1 THE PROBLEM: WHAT DO MODELS DO AND HOW WELL DO THEY DO IT?

Neuroscientists construct quantitative models to explain experimental observations of ion channels, neurons, circuits, brain regions and behavior. A model can be characterized by its *scope*: the set of observable quantities that it can generate predictions about, and by its *validity*: the extent to which its predictions agree with available experimental observations of those quantities.

Today, neuroscientists typically contribute a model to the research community by submitting a paper describing the model’s structure and scope, arguing for its novelty, and containing a very limited selection of tables and figures demonstrating its validity. Those tasked with reviewing the paper are responsible for evaluating these claims, discovering competing models and relevant data the paper did not adequately consider, and ensuring that goodness-of-fit was measured in a statistically sound manner, drawing on their knowledge of prior literature. Often, there are no means for verifying even the basic results^1^. And once a peer-reviewed modeling paper is published, it is frozen and cited as-is in perpetuity. Because publications are the sole vehicle for knowledge about model scope and validity, investigators must rely on an encyclopedic knowledge of the literature to answer *summary questions* like the following:

- Which models are capable of predicting the measurable quantities I am interested in?
- What are the best practices for evaluating the goodness-of-fit between these models and data?
- How well do these models perform, as judged by these metrics, given currently available data?
- What other measurable quantities can and can’t these models predict?
- What observations have not yet been adequately explained by any available model?

In some fields, like neuroscience, where the number of relevant publications being generated every year is growing rapidly^2^, these questions can be difficult for even conscientious senior scientists to answer comprehensively. For new computational neuroscientists, this represents a serious barrier to entry.

### 1.2 THE SOLUTION: UNIT TESTING FOR MODELS

Professional software developers face similar issues^3^. They must understand the scope of each component of a complex software project and verify that each component achieves desired input/output behavior. But software developers do not verify components by simply documenting a few interesting inputs and corresponding outputs and then presenting them to other developers for one-time review leading to an archived document. Rather, they typically follow a *test-driven development* methodology by creating a suite of executable *unit tests* that serve to specify each component’s scope and verify its implementation on an ongoing basis as it is being developed and modified^4^. Each test individually checks that a small portion of the program meets a single correctness criterion. For example, a unit test might verify that one function within the program correctly parses inputs of a certain type. Collectively, the test results serve as a summary of the project as it progresses through its development cycle. Developers can determine which features are unimplemented or buggy by examining the set of failed tests, and progress can be measured in terms of how many tests the program passes over time. This methodology is widely adopted in practice^4–6^.

Test-driven methodologies have started to see success in neuroscience as well, in the form of *modeling competitions*. During these competitions, competitors develop and parameterize models based on publicly-available training data and submit them to a central server. There, submitted models are provided hidden testing data which they must use to produce predictions. These predictions are validated using publicly available criteria to produce summaries of the relative merits of different models, just as a test suite summarizes the state of a software project. Such competitions continue to drive important advances and improve scientists’ understanding of their fields. For example, the DREAM olfaction challenge asked participants to build models predicting the odor qualities of molecules, and ranked models based on their ability to predict the odors of novel molecules^7^. The winning model in this challenge currently represents the state-of-the-art in structure-odor predictions. More relevant for neurophysiology and neuroanatomy, the quantitative single neuron modeling competition (QSNMC)^8^ investigated the complexity-accuracy trade-off among reduced models of excitable membranes; the “Hopfield” challenge tested techniques for generating neuronal network form given its function^9^; the Neural Prediction Challenge sought the best stimulus reconstructions, given neuronal activity^10^; the Diadem challenge is advancing the art of neurite reconstruction^11^; and examples from other areas of biomedical research abound^12^.

### 1.3 THE IMPLEMENTATION: SCIUNIT AND NEURONUNIT

Each of the previous examples leveraged *ad hoc* infrastructure to support model validation. While the specific criteria used to evaluate models can vary widely between modeling domains, the underlying methodology is common and therefore should be implemented once and re-used. Recognizing this, and inspired by unit testing practices, we previously developed a discipline-agnostic framework for developing *model validation test suites* called *SciUnit*^3^ (available from http://sciunit.scidash.org). *SciUnit* is now used in over a dozen scientific projects^5,6,13–29^ including The Human Brain Project^30,31^.

*SciUnit* contains validation logic common across scientific disciplines. But a particular discipline, such as neurophysiology or neuroanatomy, might have more specialized logic associated with it that can be shared amongst its sub-disciplines (e.g. hippocampal neurophysiology). We have observed a collaborative workflow where this common testing logic has been developed in common repositories on the social coding service *GitHub* (http://github.com)^6,15,18–24,27–29^. For neurophysiological and neuroanatomical tests of single neurons, we have developed such a repository, called *NeuronUnit*. This repository contains common testing logic as well as bridges to existing neuroinformatics infrastructure to make it easy to import existing datasets and models. In particular, we will describe how models described using *NeuroML*^32^ and provided freely by *NeuroML-DB*^33^ or the *Open Source Brain Project* (OSB^34^), can be validated against experimental data in a fully automated fashion, using published data curated by the NeuroElectro Project^35,36^, the Allen Institute Cell Types database^37^, or the Human Brain Project neocortical microcircuit portal^38^.

Investigators in particular sub-disciplines, for example hippocampal neurophysiologists, can use this common logic to select relevant models and tests, forming focused test suites in common repositories^20,21^. The end result of this workflow is a table like the one shown in Figure 1, where submitted models can be compared based on the scores they achieve on different selected tests. These tables serve as a summary of the status of a modeling community–just as the results from a suite of traditional unit tests serve as a summary of the status of a software project–and help scientists answer the big questions from section 1.1 more easily, thoroughly, and accurately.

**Figure 1.**
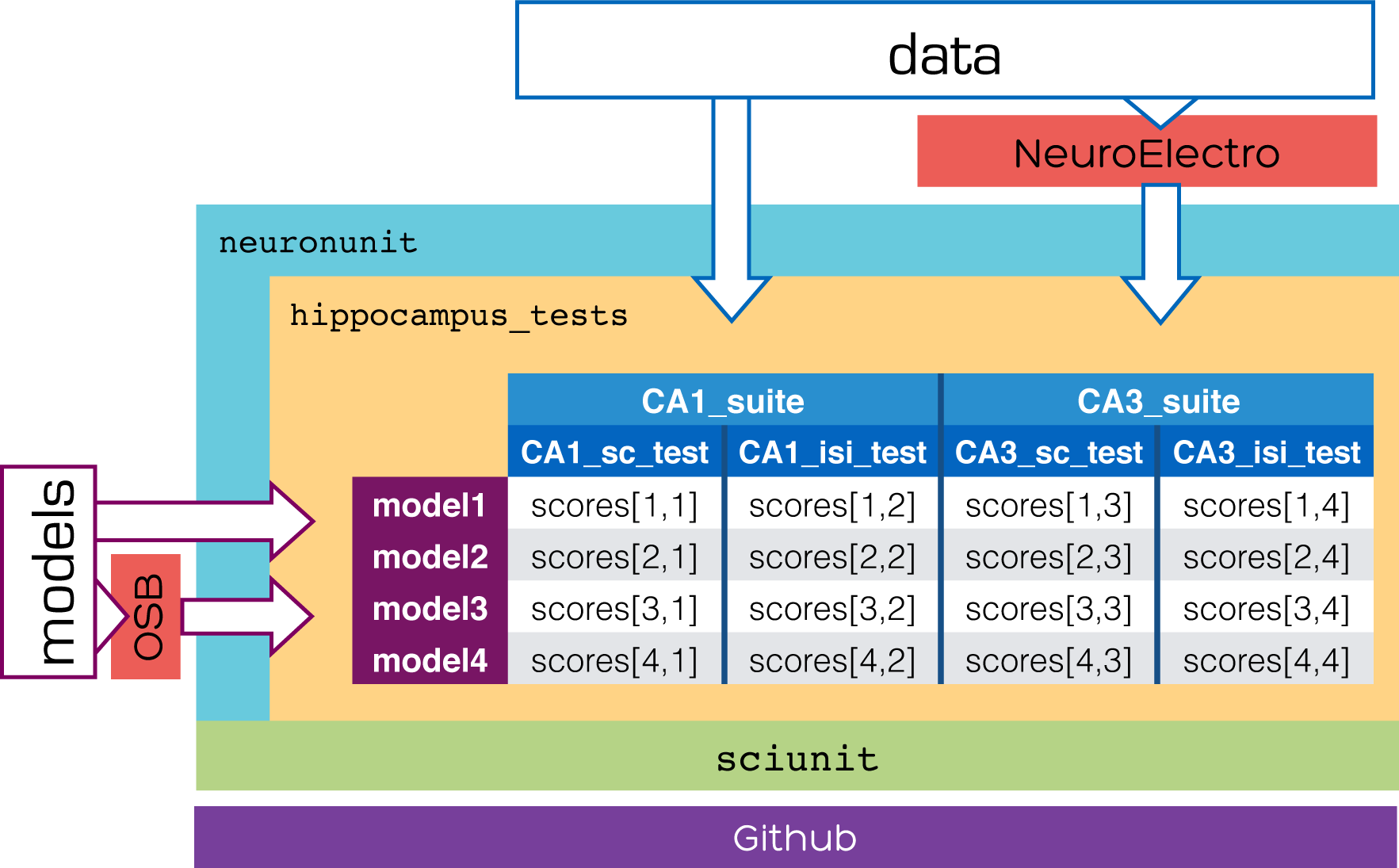
Paper overview. *NeuronUnit* is set of neurophysiology-specific testing tools built upon the domain-agnostic SciUnit framework. Scientists interested in testing neurophysiological models of particular systems, like the hippocampus, against relevant experimental data can construct test suites in a repository called, for example, *hippocampus tests*. Models and data can be added directly or imported, via *NeuronUnit*, from model repositories like NeuroML-DB or Open Source Brain (OSB) and data repositories like NeuroElectro, The Human Brain Project Microcircuit Portal, or The Allen Institute Cell Types Database. Testing tools and test repositories are developed collaboratively using Github.

## 2 VALIDATION TESTING WITH *NEURONUNIT*

### 2.1 EXAMPLE: TESTING THE RHEOBASE CURRENT

We begin by constructing a hypothetical example *SciUnit* test suite that could be used in neurophysiology. Suppose we have collected data from an experiment where current stimuli (measured in pA) are delivered to neurons of a particular subtype, while the somatic membrane potential of each stimulated cell (in mV) is recorded and stored. A model claiming to capture this neuron subtype’s membrane potential dynamics must be able to accurately predict a variety of features observed in these data. For neuron physiology, these features have been described and documented at length^36,37,39,40^.

One simple validation test would ask candidate models to predict the rheobase current, i.e. the minimum amplitude square current pulse needed to evoke at least one action potential, as shown in Fig. 2. After some agreement is reached about protocol design (how long of a square pulse, whether to do a series of fixed current steps or a binary search, what resolution to use, etc.) that protocol can be implemented (lines 15-34), and goodness-of-fit can be measured by calculating a z-score (among other possibilities) given by the difference between the predicted rheobase and the mean observed rheobase (e.g. across cells) divided by the standard deviation of the observed rheobase. The totality of one possible implementation of such a test, using *SciUnit* is shown in Figure 2. Some of the above considerations–specifically those that are not specific to the problem domain such as calculation of a z-score–are automatically handled by the internal logic of *SciUnit* through the `sciunit.Test` class and thus are not shown explicitly. Importantly, in contrast to the simple example in Fig. 2, the actual code for the `RheobaseTest` in *NeuronUnit* contains additional logic for speed, handling edges cases and exceptions, using existing standards, and other considerations. However, the code shown in Fig. 2, lines 1-34 is a good representation of the core idea.

**Figure 2.**
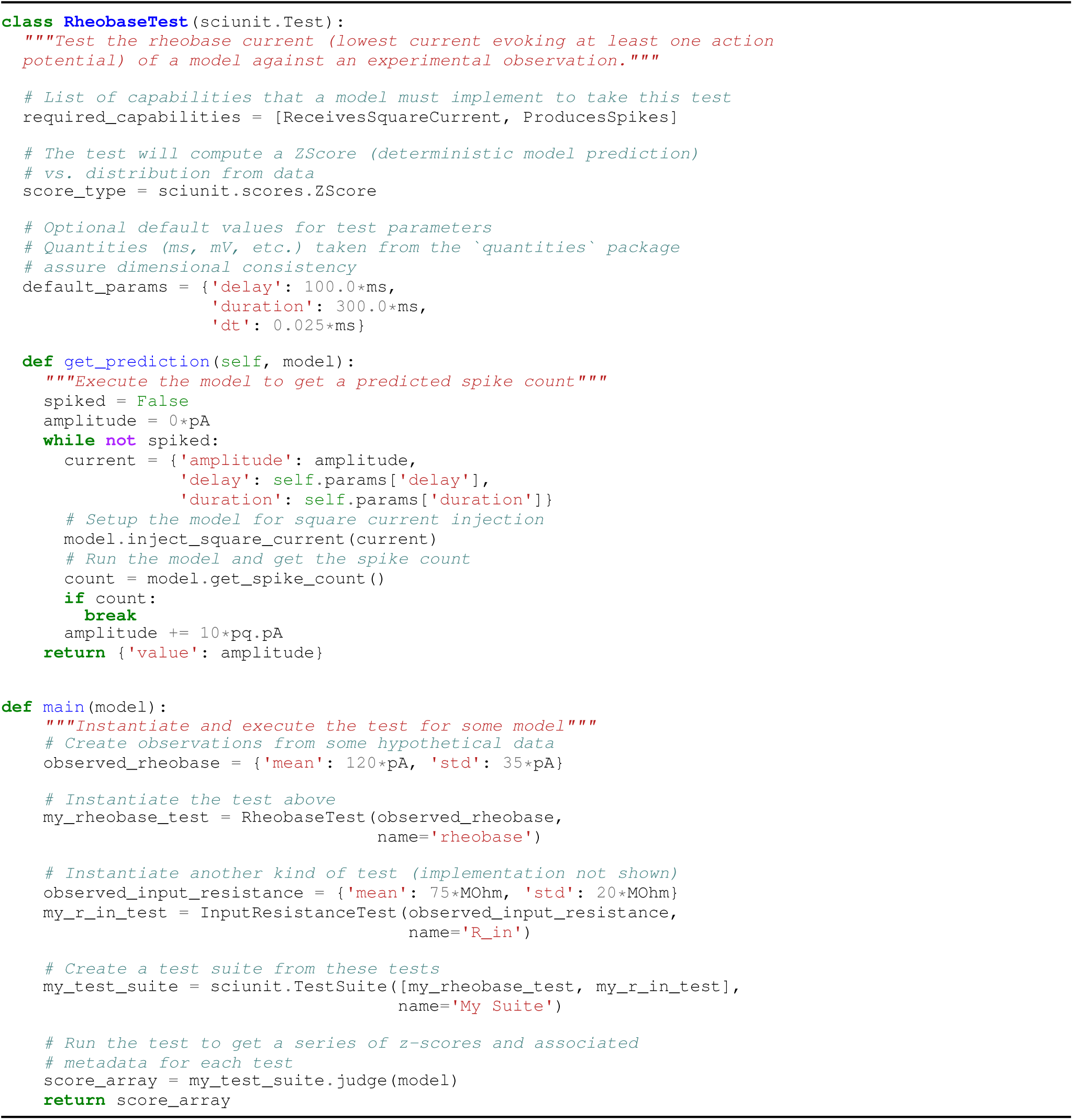
A toy example of a single neuron rheobase test class implemented *directly* using *SciUnit* without *NeuronUnit*. Because this implementation contains logic common to many different systems, *NeuronUnit* was developed to provide a simpler and more robust means to deliver it.

### 2.2 CONSTRUCTING AND EXECUTING VALIDATION TESTS IN *NEURONUNIT*

A *SciUnit* validation test is an instance (i.e. an object) of a Python class implementing the `sciunit.Test` interface (Fig. 2, line 1). In order to instantiate it, we must at least provide the observation that parameterizes it (line 40). We emphasize the crucial distinction between the *class* `RheobaseTest`, which defines a *family* of possible validation tests but is not yet tied to any particular experiment observation, and the particular *instance* `my_rheobase_test` defined on lines 43-44, an individual validation test parameterized by a particular set of observed experimental data, such as from cells recorded in one investigator’s lab.

Alongside this *rheobase test*, we might also specify a number of other tests capturing different features of the data to produce a more comprehensive suite. For example, the QSNMC defined 17 other validation criteria in addition to one based on the overall spike count, capturing features like spike latencies (SL), mean subthreshold voltage (SV), interspike intervals (ISI) and interspike minima (ISM) that can be extracted from the data given appropriate stimulation protocols^8^. On lines 47-48 we create the observation and test corresponding to one other possible criterion: the input resistance of the cell. The QSNMC authors defined a combined metric favoring models that broadly succeeded at meeting these criteria, to produce an overall ranking. Such combinations of tests, including any weights assigned to each, constitute a *test suite*, handled by `sciunit.TestSuite`, as shown in lines 52-53.

Communities have built *SciUnit* test repositories capturing the criteria used in their subfields of neuroscience^6,15,18–24,27–29^. Consequently, test generation for a particular system of interest often requires simply instantiating a previously-developed class with particular experimental parameters and data. We developed *NeuronUnit* (http://github.com/scidash/neuronunit) to define many such classes of models and tests for neurophysiology and neuroanatomy. In contrast to the toy example in Figure 2 used for illustration, these tests are thoroughly unit-tested (in the traditional software development sense) to handle alternative parameter specifications, edge cases, numerical issues, etc. *NeuronUnit* currently implements nearly 50 such test classes. Particular instantiated tests, such as a hypothetical `CA1_rheobase_test`, can then be derived from these classes in suite repositories (e.g. `hippocampus_tests`, as shown in Figure 1).

### 2.3 CAPABILITIES THROUGH ELEPHANT

All test classes, by virtue of inheriting from `sciunit.Test`, must contain a `get_prediction` method (Fig. 2, line 19) that receives a candidate *model* as input and produces a *prediction* dictionary as output. Other methods such as `compute_score` can be provided, or they can default to the methods associated with the parent test class or the provided score type. To specify the interface between the test and the model, the test author provides a list of *capabilities* in the `required_capabilities` attribute (Fig. 2, line 6). Capabilities are simply classes describing methods that a test needs a model to implement so that the test can provide inputs and generate relevant predictions using the model. They are analogous to *interfaces* in e.g. Java. In Python, capabilities are written as classes with unimplemented members. The *NeuronUnit* capabilities required by the test in Fig. 2 are shown in truncated form in Fig. 3. The test in Figure 2 uses specific methods of these capability in lines 28 and 30 to produce a spike count prediction for each input current. When a model cannot take a test because of a missing capability, a `NotApplicable` score is returned (e.g. spiking neuron models describing governed entirely by point processes cannot predict subthreshold voltages) when the test is executed. Thus models which are a poor match to the data can be distinguished from models which cannot even in principle match the data.

**Figure 3.**
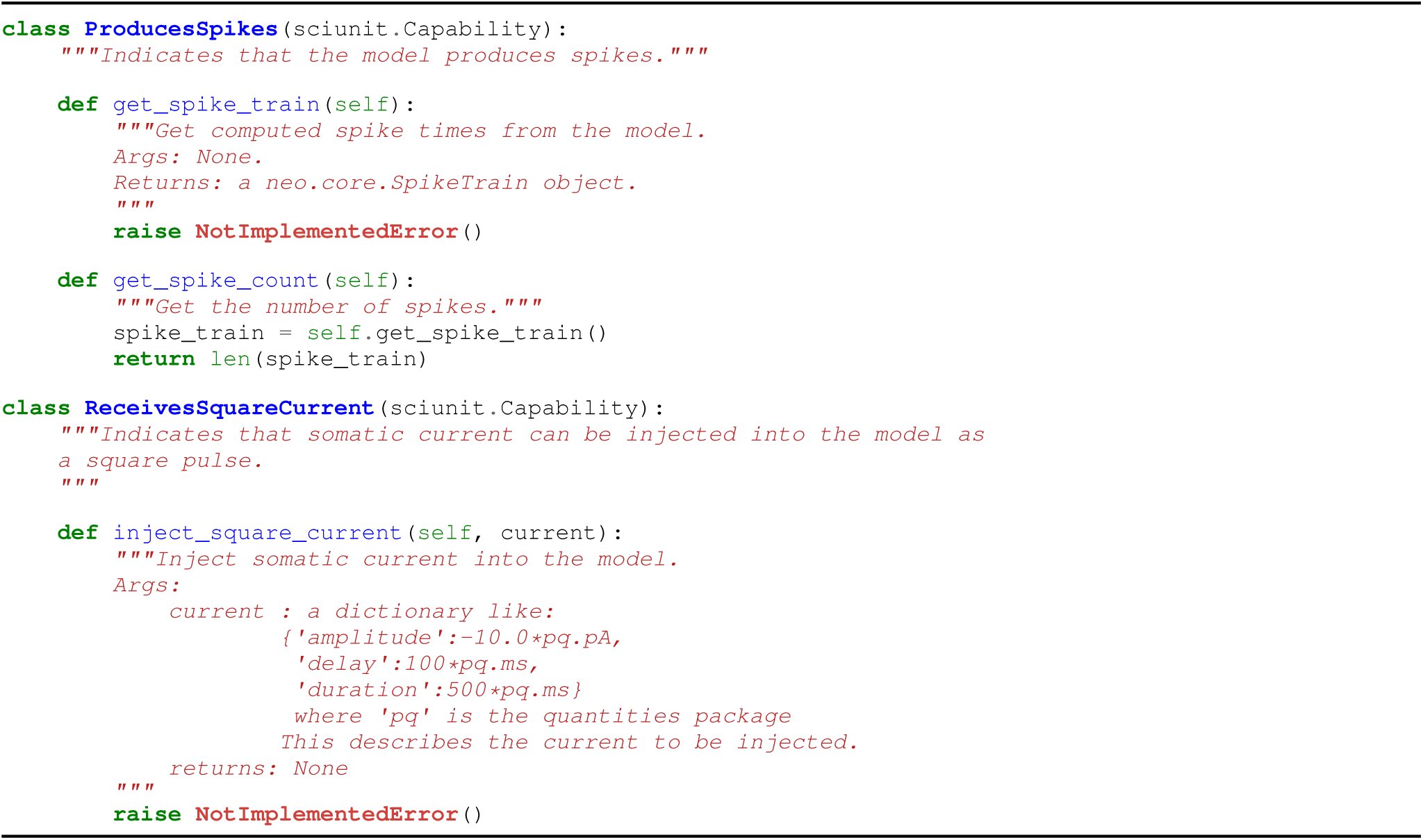
Truncated versions of two capability classes describing the methods (and inputs and outputs) that a model must implement in order to take a test that requires either the ProducesMembranePotential or ReceivesCurrent capability (or both) of the models that it judges. All methods in each capability must have an implementation for the model to satisfy the requirement, but some (see `get_spike_count`) are automatically implemented by the implementation of others (see `get_spike_train`). *NeuronUnit* defines many such capabilities, and separately some reference implementations of those capabilities using various simulator backends including NEURON in conjunction with the Elephant analysis library to, for example, extract a spike train from a membrane potential time series, or a membrane potential time series from a model simulation.

Elephant^41^ is a Python library supporting tasks associated with analysis of neural data (or model output), such as membrane potential time series, spike trains, etc. as described by the neuron electrophysiology object library Neo^42^. It is an open source and actively developed project, containing reliable algorithms on which to base neurophysiology tests. *NeuronUnit* uses Elephant to give models a path for implementing a variety of capabilities used by *NeuronUnit* tests. For example, the *NeuronUnit* capability `HasSpikeTrain` requires that a method named `get_spike_train` be implemented, which means that it must (by whatever means) return a Neo `SpikeTrain` object corresponding to the train of spikes produced by the model under some conditions, e.g. in response to a certain stimulus. `NeuronUnit` models aiming to implement this capability can do so in part by calling the Elephant method `AnalogSignal.threshold_detection`, which takes a Neo `AnalogSignal` object (e.g. a membrane potential time series) and returns a Neo `SpikeTrain` object (Fig. 4, lines 20-26). This reduces the problem of generating the spike train to one of obtaining a membrane potential time series, which might plausibly vary across models. The membrane potential can be obtained by calling the `get_membrane_potential` method of the `HasMembranePotential` capability, with a somewhat more complex and nested implementation that also relies on Neo objects (Fig. 4, lines 8-18). Elephant is used in this manner to implement a variety of *NeuronUnit* capabilities for *NeuronUnit* model classes, without requiring that the modeler do so explicitly (though they can override these if they wish in their own model class definitions). This also helps neuroscientists by directing them to a set of common utility functions (in Elephant) and data object types (in Neo) that they can use, knowing that they are well-tested and accepted by the community.

**Figure 4.**
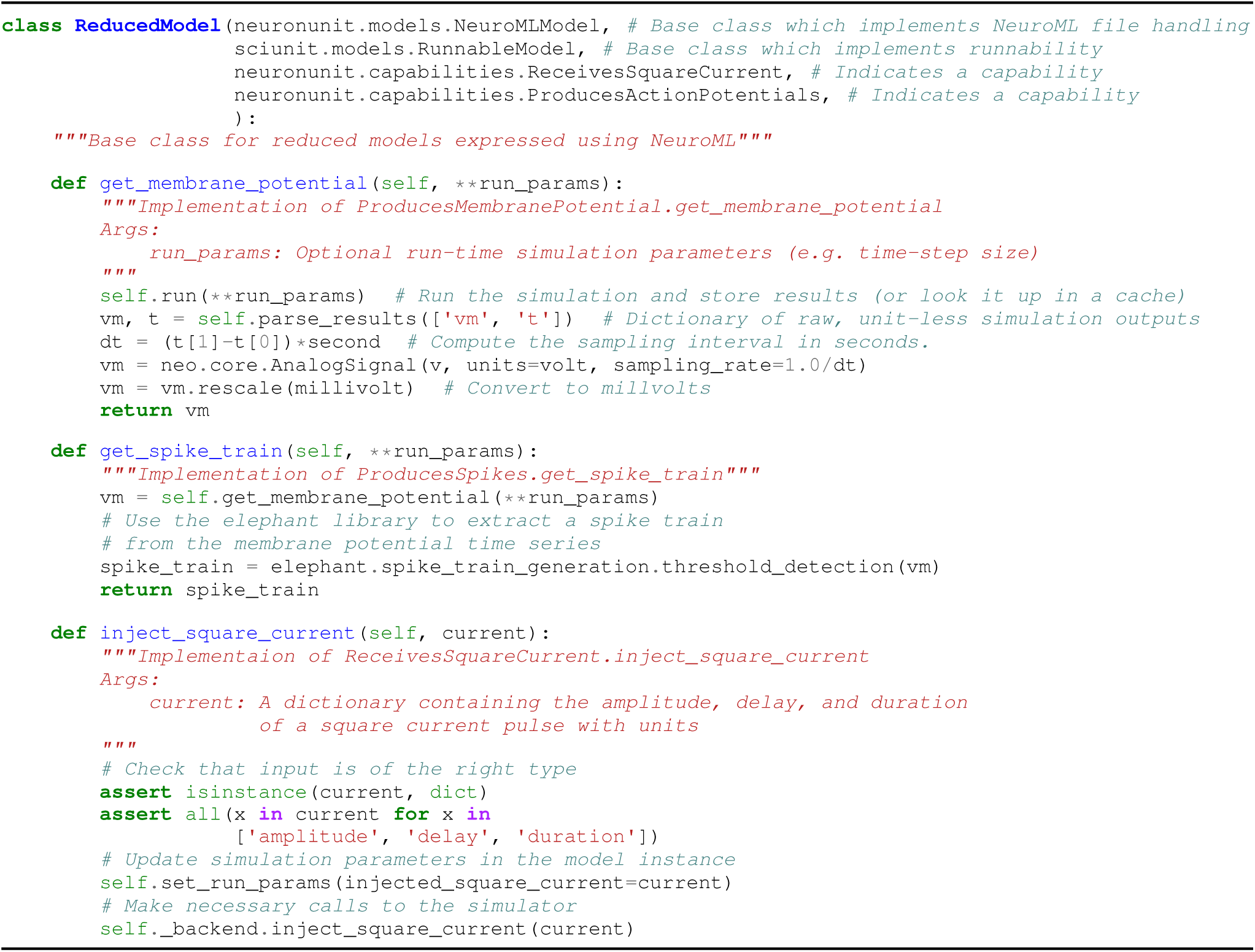
A class for reduced models. This class provides simple implementations of the capabilities required by the *NeuronUnit* test outlined in Fig. 2, and other tests requiring such capabilities. The constructor, which is inherited from `NeuroMLModel` and thus not shown, takes a file path or URL to a NeuroML file, and an optional simulator backend.

In *SciUnit*, the `judge` method (Fig. 2, line 57) checks that these capabilities are implemented by the model, uses them to extract a prediction via *get prediction* (Fig. 2, lines 19-34), to compute a score indicating model/experiment agreement (not shown) and return a score. In the example from Fig. 2 a z-score was specified (line 10), so an instance of `sciunit.scores.ZScore` is computed and returned. In addition to the score itself, the returned score object may also contain metadata, via the `related_data` parameter, for those who may wish to examine the result in more detail later.

### 2.4 MODELS IN NEUROML

Capabilities are *implemented* by models. In *SciUnit*, models are instances of Python classes that inherit from `sciunit.Model`. Like tests, the class itself represents a family of possible models. A particular model is an instance of such a class, parameterized by the arguments of the constructor

NeuroML is a standardized model description language for neuroscience^32^. It allows many neurophysiological and neuroanatomical models to be described in a simulator-independent fashion. Some simulators, notably the popular NEURON package^43^, can seamlessly export model specifications as NeuroML. Because NeuroML is an XML specification, model descriptions can be verified for correctness and queried for arbitrary information. The latter facilities allow model parameters and capabilities to be automatically determined and also mapped to appropriate variables or functions in simulators. A number of popular simulators support execution of NeuroML models, including NEURON, GENESIS^44^, NEST^45^, and MOOSE^46^. Additionally, over 1500 cell models spanning dozens of cell types can be obtained in NeuroML format from NeuroML-DB (http://neuroml-db.org).

To take advantage of the transparency and machine-readability of models described using NeuroML, *NeuronUnit* provides a `sciunit.Model` subclass called `NeuroMLModel`. It can be instantiated directly or sub-classed (useful when additional capabilities need to be manually exposed, for example) to generate a model suitable for testing (Fig. 4). One creates such an instance of a `NeuroMLModel` by passing the path or URL to a NeuroML model file to its constructor.

Figure 4 shows a truncated version of such a model class which can support the execution of models described by NeuroML files (by inheriting from `neuronunit.models.NeuroMLModel` and which implements the capabilities described in Fig. 3. The implementation of capabilities may rely on third party libraries which have been established for specific tasks in the analysis of neural data (real or simulated). For example, as described above the conversion of a simulated membrane potential time series to a discrete spike train is handled by a function from the Elephant library^41^. Altogether, this model is sufficient to run any test requiring (at most) those capabilities, such as the rheobase test in Fig. 2 or a number of other tests of the features or dynamics of membrane potential time series or spike trains.

### 2.5 SIMULATOR BACKENDS AND CACHING

Simulation itself is handled via one of several simulator `Backend` classes. These backends implement a bridge between the model class and the simulator. In some cases, the backend may simply use existing Python bindings provided by the developers of the simulator. The backend is selected when the model is instantiated (Fig. 4, see caption). In principle, simulating the same conceptual model (e.g. a single NeuroML file) using different simulator backends may produce the same results, and therefore the same test scores, although this cannot be guaranteed. Furthermore, there may be no adequate route for simulating the model described by a NeuroML file on a given simulator, in which case a backend for a more appropriate simulator should be selected.

NeuronUnit currently provides backends for (1) jNeuroML, a NeuroML reference simulator and (2) NEURON, the most widely used simulator for neuron models. Other backends are in development. PyNN is a python modeling library for networks of simple neurons with both broader and deeper coverage of simulators than NeuronUnit offers^47^. Therefore one potential route to broader coverage of simulators in network models would be to create a backend which wraps model interactions in PyNN calls, re-using PyNN’s simulator communication infrastructure.

The base `Backend` class supports caching, such that repeated calls to run the same simulation under the same parameters will simply looks up simulation outputs in the cache rather than unnecessarily run the simulation again. This frequently occurs when different tests require the same simulation outputs in order to compute features. For example, a test of the width of an action potential requires the same simulation output as the test of the height of an action potential. Therefore only the first of these tests causes the simulation to be run. The second of these tests will operate on cached simulation output to compute a different feature.

### 2.6 TEST SUITES AND SCORE MATRICES

A collection of tests intended to be run on the same model can be put together to form a test suite, as in Fig. 2, lines 51-53. Like a single test, a test suite is capable of judging one or more models. To make the idea of a test suite more concrete, we created a test suite from six *NeuronUnit* tests that compare basic electrophysiological properties of model neurons to their experimental counterparts. Specifically, we instantiated these tests using data for olfactory bulb mitral cells taken from http://neuroelectro.org^35,36^. We then executed this test suite against seven published models of olfactory bulb mitral cells. The output of this test execution is a `ScoreMatrix` containing a score for each model on each test (Fig. 5). A `ScoreMatrix` is a subclass of a Pandas dataframe^48^ through which scores for a given model or a test can be summarized, visualized, or probed for further details. Each row or column of a `ScoreMatrix` is a `ScoreArray` (a subclass of a Pandas series) summarizing the scores for one model or test, as returned in line 57 of Fig. 2.

**Figure 5.**
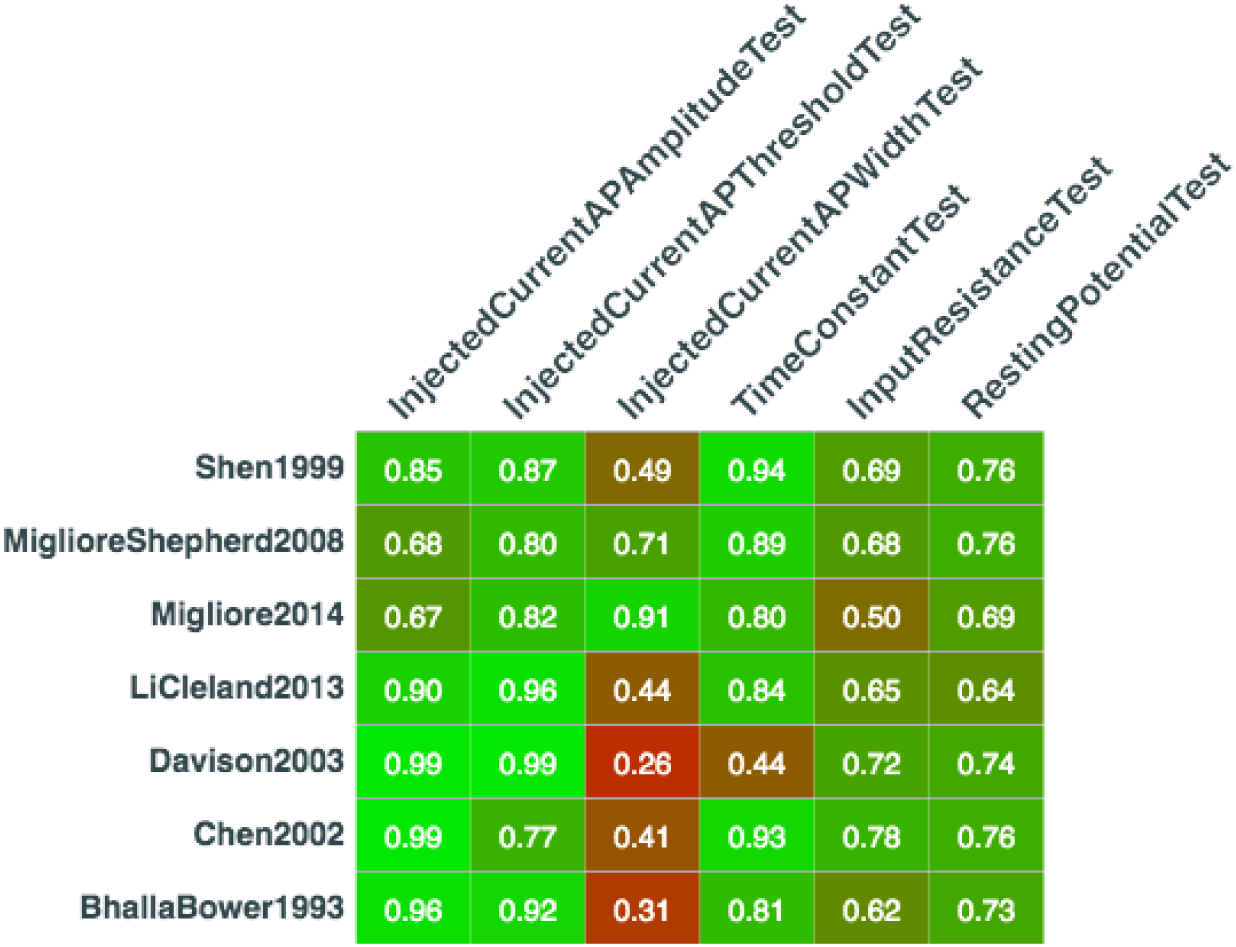
A score matrix for a SciUnit test suite, visualized as a hyperlinked table inside a Jupyter notebook stored in a test repository at http://github.com/justasb/mitralsuite. Individual elements of this score matrix can be further probed to see test and model details as well as simulation outputs or experimental data that gave rise to these scores. For ease of visualization, each raw score (a z-score comparing model output to the experimental data distribution) has been automatically converted into a normalized score ranging from 0 (lowest possible agreement) to 1 (best possible agreement, i.e. that feature of the model’s output is equal to the mean of the same feature in the experimental data distribution).

### 2.7 TRAINING AND OPTIMIZATION

Some models could in principle match experimental observations, but not under the current set of parameters. Optimizing a model’s parameters through testing could produce a model which is a better description of the underlying system. We have been developing test-driven model optimization in *NeuronUnit* using a combination of Scoop^49^ (for parallelism), DEAP^50^ (for genetic optimization), and BluePyOpt^51^. This allows models to be optimized against suites of *NeuronUnit* tests, either weighting the tests to produce “best” models for the suite, or retaining the entire non-dominated set, i.e. all model parameterizations that are not strictly worse than some other parameterization on every test. This latter option may be important to avoid producing models which have the greatest correspondence to the *mean* value of all experimental data features for a neuron type, but which do not correspond to the behavior of any particular neuron^52^. We have used this feature internally to optimize reduced neuron models against suites of tests of electrophysiological properties in order to produce sets of reduced models for each of several major cell types in the brain. We plan to release these optimized models as“plug and play” options for investigators who wish to build circuit models containing several distinct, realistic, neuron subtypes. However, this features is not yet production ready.

### 2.8 MODELS IN OPEN SOURCE BRAIN

Open Source Brain (*OSB*^34^) curates many models described in NeuroML, and is a hub for collaborative modeling. In constrast to NeuroML-DB, OSB is intended to be a platform for on-going model development. OSB-curated projects have been converted from their native format (e.g. NEURON) into NeuroML. To ensure that the NeuroML model is faithful to the original simulation, OSB verifies concordance between model output (beginning with the NeuroML description) and reference output (from native simulator source files or published figures) for each model. Thus, OSB is an excellent source of modeling projects that, in addition to being open source, are known to be accurately described. This also highlights the distinction between *verification* activities, performed by OSB to ensure that a model description operates correctly, and *validation* activities, performed using *NeuronUnit* to check the model’s predictions against data. Both are important activities. *NeuronUnit* also implements an `OSBModel` class for those who wish to start testing an active modeling project on OSB.

### 2.9 MORPHOLOGY TESTS

Some neuron models aim to have realistic neuron morphologies. *NeuronUnit* provides a set of morphology tests that use pyLMeasure^53^, a python interface to the LMeasure^54^ tool for generating morphological features from both model neurons and reconstructions of real neurons. These tests can be parameterized with either features of some real neuron’s morphology, or distributions of such features across ensembles of real neurons. In order to take these tests, models must implement a `ProducesSWC` capability, which means the model class must have a means of generating a standard .swc file (as used in neurite reconstructions) corresponding to its morphology. For models expressed in NeuroML or NEURON, this is an easy capability to implement.

## 3 DATA SOURCES

In the examples in Fig. 2, tests were provided data explicitly (e.g. lines 40 and 47). However, experimental data about neurons are also available from existing community resources in standardized formats. In this section, we will show how to use this existing infrastructure to parameterize tests with published data automatically.

### 3.1 NEUROELECTRO

The NeuroElectro project (http://neuroelectro.org) is an effort to make all published data on single cell neurophysiology available in a machine-readable format^55^. Currently, up to 47 electrophysiological properties are available for 235 cell types, gathered from over 2000 single pieces of published data extracted from tables in published papers. NeuroElectro uses the NeuroLex.org ontology^56^ to identify specific neuron types and specific electrophysiological properties, and indexes corresponding data values from the literature. We have made it easy to parameterize the test families in *NeuronUnit* using data extracted from the NeuroElectro API. Tests can be based upon data from single journal articles, or from ensembles of articles with a common theme (e.g. about a particular neuron type). The latter is illustrated in Figure 6. Once data has been retrieved from NeuroElectro, associated statistics (e.g. mean, standard error and sample size) are extracted to parameterize tests (e.g. a test of a model cell’s input resistance). Data from NeuroElectro can be used to validate a variety of basic electrophysiological features of neuron models, including spike width, resting membrane potential, after-hyperpolarization amplitude, etc. As NeuroElectro is the only public, curated source of such data for many neuron types, it represents a key resource facilitating test construction. However, in selecting data from NeuroElectro (either averaged data across all reports, or data from selected reports) for test construction, one is implicitly assuming that it represents the kind data that ought to be binding in model development. For some models, data obtained in this way ought not to guide model development, and other data sources will need to be identified and used for test construction.

**Figure 6.**
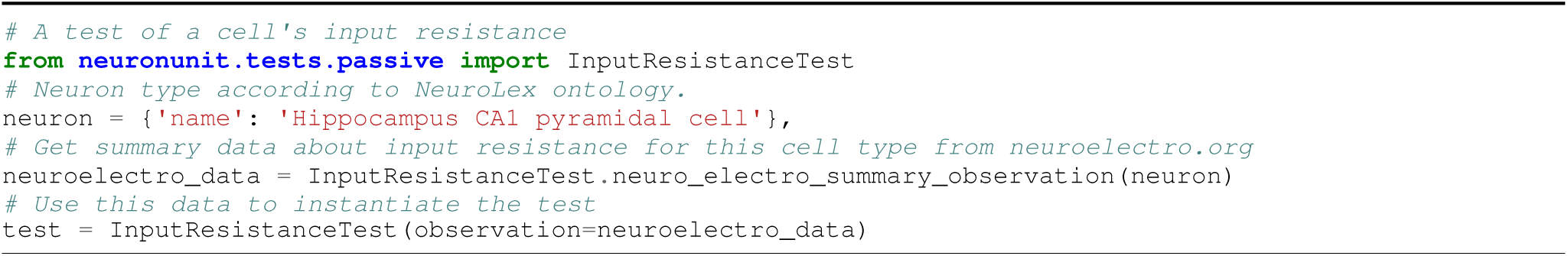
Tests can be built directly from cell-type-specific data automatically pulled from http://neuroelectro.org.

### 3.2 OTHER PUBLIC DATA SOURCES

For those interested primarily in neurons from either visual cortex or somatosensory cortex, *NeuronUnit* can also automatically build tests using data from The Allen Institute’s Cell Types Database^37^ and The Human Brain Project’s Neocortical Microcircuit Collaboration Portal^38^, respectively. *Neuronunit*’s `bbp` and `aibs` modules provide the means to communicate with the corresponding APIs or download data directly, to extract recordings that can be used to compute the features used in observations that will parameterize a *NeuronUnit* test.

## 4 DISCUSSION

Neuroscientists can use *NeuronUnit* to easily create test suites tailored to specific neuronal systems of interest. For example, investigators interested specifically in testing models of cells in the hippocampus would use *NeuronUnit* to create tests in a GitHub repository called e.g. *hippocampus tests*, containing scripts or Jupyter notebooks containing the steps needed to instantiate and execute validation tests parameterized by empirical data from the hippocampus against corresponding hippocampal cell models. Many of the other features and details of *NeuronUnit*, including specific examples for developers, can be found at http://github.com/scidash/neuronunit.

### 4.1 DIFFERENCES OF OPINION

Validation criteria are subject to debate (indeed, the QSNMC criteria changed between 2007 and 2008 due to such debates), and identification and correction of flawed methodology is becoming increasingly common^57–59^. Statisticians and other scientists interested in improving how goodness-of-fit is measured can simply fork the *NeuronUnit* repository, modify the logic in the relevant test families and submit a *pull request* for evaluation by the community. Rather than attempting to persuade a large number of scientists that the methods used in canonical papers need to be changed, statisticians need only propagate a change to the testing logic used by the community. Once the testing logic has been changed, all models can be immediately evaluated against the new metrics. Those who wish to identify statistical best-practices for measuring goodness-of-fit need only refer to the tests in these repositories.

### 4.2 COMMUNITY CONTRIBUTIONS

To submit a new model for testing, one can similarly fork a test repository (e.g. *hippocampus tests*), add a new model and test it locally using the tests in the repository. When satisfied with parameter choices, one can submit a pull request to the upstream repository. Once a model is in the repository, it can be evaluated on an ongoing basis against the latest accepted collections of tests, parameterized by the latest data, using the latest statistical best practices, without the participation of the original modeler, assuming only that the capabilities required by the tests stay stable.

GitHub is an excellent platform for this kind of activity, and is becoming a widely-used tool for scientific collaboration, reproducibility, and transparency^60^. We anticipate that other communities within neuroscience (and indeed, within science more broadly) will develop repositories similar to *NeuronUnit* that provide a common, collaboratively-developed vocabulary of test families and capabilities relevant to their domain, and bridge relevant standardization efforts and infrastructure. Using these common repositories, smaller groups of investigators interested in particular systems of interest will develop tests and models in test repositories like *hippocampus tests*. Each research community can develop suitable quality standards and processes for merging pull requests.

### 4.3 PUBLIC ASSESSMENT

The most visible result of these efforts will be tables that allow investigators to represent, formally, the state of model development in their field, as judged against corresponding experimental data. The `ScoreMatrix` is an implementation of such a table, and when rendered in a Jupyter notebook it appears similar to the one shown in Figure 5. Thus a Jupyter notebook, in addition to representing a reproducible workflow for testing, can also be used to visualize the test outputs themselves.

We are also developing a lightweight portal, *SciDash*, that directs investigators to the canonical tests in their field in order to prevent fragmentation and encourage community participation. *SciDash* will also facilitate viewing test results without needing to clone test repositories and run tests locally for exploratory purposes. Finally, *SciDash* will support the scheduling and execution of tests for those who prefer a web interface to a notebook or IDE. *SciDash* will not be a place for the *development* of models, as existing services such as OSB and NetPyne^61^ are already well-suited to that purpose.

### 4.4 MANUALLY CREATING NEW MODELS, CAPABILITIES, AND TESTS

*NeuronUnit* provides base classes to enable rapid generation of models, capabilities, and tests for neurophysiology and neuron morphology data. However these objects can also be created from scratch, requiring only adherence to the *SciUnit* interface as exemplified in sec. 2. For example, a Model could implement an `integrate` capability method by wrapping execution of a MATLAB script and a `get_spikes` capability method by parsing a resulting .csv file on disk; a Test could be initialized using empirical spike rates collected in the lab. While this does not meet our idealized vision of collaborative development and testing, in practice this may be a common scenario.

### 4.5 PARTICIPATION FROM MODELING COMMUNITIES

Modelers may not want to formally expose their models to tests via capabilities, a requirement for test-taking. We anticipate four solutions: **First**, we will emphasize to the community that wrapping existing model code that satisfy a capability is quite simple, requiring as little as one line of code. Importantly, the modeler is not required to expose or rewrite any internal model flow control in almost all cases. **Second**, we support multiple simulation environments automatically by using NeuroML^32^, and other simulator-independent model descriptions are possible for other domains. Automated generation of NeuroML from native model source code is in development (Gleeson, personal communication); for the popular NEURON simulator^43^, this functionality is already implemented and in use for some model components. This minimizes modeler effort for a large (at least 1500) and growing number of neuronal models that are not already in NeuroML-DB. However, models converted from native simulator code to NeuroML must still be examined manually to ensure that nothing has been lost in translation. **Third**, modelers have an incentive to demonstrate publicly their models’ validity. Participation in public modeling competitions demonstrates that this incentive is sufficient to recruit several participants. We plan to assist with such competitions by advocating for use of *SciUnit* as the underlying technical substrate in the future. Currently, we are in discussion with field specialists to organize competitions for spiking neuron models (a sequel to the QSNMC), spike-sorting algorithms, calcium imaging spike time reconstruction, brain-computer interfaces, neural network function prediction, and seizure prediction. **Fourth**, modelers have an incentive to use *SciUnit* or *NeuronUnit* during development (see TDD, Sec. 1) to ensure that ongoing development preserves correspondence between model and data. A popular test suite can represent a “gold standard” by which progress during development is judged, even within a single project.

### 4.6 PARTICIPATION FROM EXPERIMENTAL COMMUNITIES

Experimentalists may not want to write tests derived from their data. We anticipate four solutions: **First**, we will emphasize that tests do not require releasing entire datasets; a test consists only of a list of required capabilities (for selecting eligible models), and sufficient statistics suitable for executing the scoring logic. This can simply be means and standard deviations in many cases, such as the examples in this paper. Most scoring logic will consist of a few calls to capability methods followed by a standard statistical test, which can be implemented once in *SciUnit* or a discipline-specific repository like *NeuronUnit*. **Second**, data-sharing is becoming accepted. If data is available from a community repository, as many granting agencies are increasingly requiring, then tests can often be generated automatically using bridges developed collaboratively by the community. **Third**, an incentive to write tests for one’s data exists: the ability to identify models that give the data clear context and impact. If a new piece of data invalidates a widely-accepted model, this can be cited in publications. **Fourth**, one can compare new data against existing data by simply creating a *data-derived model* that directly generates “predictions” by draws from the existing dataset directly. Thus the model validation procedures described in this paper can also be used to perform data validation – that is, check the goodness-of-fit of new data with old data.

### 4.7 OCCAM’S RAZOR

All things being equal, simpler models are better. Model complexity, though fundamentally a subjective measure, has many correlates^62^, including: 1) model length; 2) memory use; 3) CPU load; 4) # of capabilities. Python packages for reporting these complexity measures exist^63^, and can be used in conjunction with test scores themselves to visualize the model validity vs complexity tradeoff, for example as a scatter plot, with the “best” models being in the high validity / low complexity corner of the plot. The set of models which *dominate* all others, i.e. that have the highest validity for a given complexity, can be represented as a “frontier” in such a scatter plot (e.g. as used in the symbolic regression package Eureqa^64^).

### 4.8 EXPANSION INTO OTHER AREAS OF NEUROSCIENCE

*NeuronUnit* largely covers the neurophysiology and morphology of single neurons. We would like to extend both it and the larger *SciUnit* effort across neuroscience and the other biological sciences. The *SciUnit* framework is discipline-agnostic, so community participation and model description are the only obstacles. Community participation begins with enumerating the capabilities relevant to a sub-discipline, and then writing tests. Model description can expand within NeuroML (which already covers multiple levels within neuroscience) and tools for capability implementation can incorporate libraries for neuroimaging (NiBabel, http://nipy.org/nibabel), neuroanatomy (NeuroHDF, http://neurohdf.readthedocs.org) and other sub-disciplines. SBML will enable expansion beyond neuroscience, facilitated by parallel efforts among NeuroML developers to interface with it (Crook, unpublished). *SciUnit* has already been used extensively at several levels from ion channels to behavior in the OpenWorm project^5,6,17^, which through open, collaborative development seeks to model the entire organism *C. elegans*.

### 4.9 PROJECT DEVELOPMENT

Gewaltig and Cannon describe the maturity of a computational software project^65^ as “review ready”, “research ready”, or “user ready”. The development of the underlying *SciUnit* framework is complete^3^, and it provides the full interface necessary for continued development of domain-specific unit-testing frameworks – it is research ready if not user ready. *NeuronUnit* itself is “review ready” – we invite others to participate in building it out to satisfy the use cases they encounter. Meanwhile, corresponding test suite repositories on GitHub, containing *NeuronUnit*-based tests, do not even reach this stage (with few exceptions^21^). However, many neuroscience projects are realizing this vision using *SciUnit* directly^5,6,15–20,24–29^. We plan to continue creating such repositories; more importantly we invite interested parties to create their own, either to guide in-house model development or to assess larger pools of established models.

## ACKNOWLEDGEMENTS

We thank Jonathan Aldrich, Shreejoy Tripathy, and Padraig Gleeson, and MetaCell LLC for their support of this project. The work was supported in part by grant R01MH081905 and R01MH106674 from the National Institute of Mental Health, and grant R01MH106674 from the National Institute for Biomedical Imaging and Bioengineering. The content is solely the responsibility of the authors and does not necessarily represent the official views of the National Institutes of Health.

